# Genetic analyses of medication-use and implications for precision medicine

**DOI:** 10.1101/501049

**Authors:** Yeda Wu, Enda M. Byrne, Zhili Zheng, Kathryn E. Kemper, Loic Yengo, Andrew J. Mallett, Jian Yang, Peter M. Visscher, Naomi R. Wray

## Abstract

It is common that one medication is prescribed for several indications, and conversely that several medications are prescribed for the same indication, suggesting a complex biological network for disease risk and its relationship with pharmacological function. Genome-wide association studies (GWASs) of medication-use may contribute to understanding of disease etiology, generation of new leads relevant for drug discovery and quantify prospects for precision medicine. We conducted GWAS to profile self-reported medication-use from 23 categories in approximately 320,000 individuals from the UK Biobank. A total of 505 independent genetic loci that met stringent criteria for statistical significance were identified. We investigated the implications of these GWAS findings in relation to biological mechanism, drug target identification and genetic risk stratification of disease. Amongst the medication-associated genes were 16 known therapeutic-effect target genes for medications from 9 categories.

## Introduction

Susceptibility to most common human diseases is complex and multifactorial, involving both genetic, environmental and stochastic factors^1^. During the last decade, large-scale genome-wide association studies (GWASs) have identified thousands of single nucleotide polymorphisms (SNPs) associated with diseases and related traits, consistent with a polygenic genetic architecture of common disease. These results add useful human-relevant information to drug development, drug repurposing and clinical trial pipelines^2^. Here, we have turned the tables, aiming to identify genetic loci associated with medication-taking. In the context of electronic health record data, medication-use may be an easy route to identify disease-case subjects. However, in clinical practice, it is common that one medication is prescribed for several indications, but conversely, several medications can be prescribed for the same indication. It is likely that medication-use reflects not only similarity between different clinical manifestations^3^ and/or comorbidity^4^ of diseases but also heterogeneity of clinical manifestation (symptoms and signs) and of intervention response (for example, from lifestyle change to the combination of treatments).

We hypothesise that genetic variants associated with taking medications categorised based on anatomical and therapeutic classifications may add additional relevant information to understanding the underlying biological mechanism of diseases and drug development approaches. Here, we study genetic variation in current medication-use. We report 505 loci independently associated with medication categories. We explore these GWAS findings for biological mechanisms and as drug targets. We estimate the genetic correlation between the 23 medication traits, and with other diseases and traits using published GWAS results. We use Mendelian Randomization to investigate putative causal relationships among diseases and traits. We show that genetic predisposition to common disease predicts likelihood of taking relevant medications, a significant finding in relation to future practice of precision medicine for common disease.

## Results

### Case-control GWAS of medication-use

Medications taken by UKB participants were classified using the Anatomical Therapeutic Chemical Classification System^5^ and provided in **Figure 1** and **S1**, **Table S1**. **Figure S2** shows the demographics of participants with medication records. The full phenotype extraction pipeline for UKB participants is summarised in **Figure S3**. An overview of analyses is provided in **Figure S4**. The medication-use case-control GWASs identify 910 within-trait independent SNPs significantly associated (P < 5×10^−8^) across 23 medication traits (**Figure 2 and S5**). After applying a more stringent multiple testing threshold (P < 1e-8/23) ^6^, a total of 505 SNPs remain (**Table S2 and S3**), with per-trait associations ranging from 0 (C02: hypertensives, N02A:opioids, N06A: antidepressants) to 103 (C09: agents acting on renin-angiotensin system) SNPs. Many of the associated SNPs may simply be a reflection of the primary indication for which the medication is prescribed (**Table S4**). For example, C09 medications have therapeutic effect on hypertension; of the 103 independent SNPs associated with C09 medications (P < 10^−8^/23), we identified SNPs previously linked to hypertension (7 SNPs)^7^, systolic blood pressure (32 SNPs)^8^, diastolic blood pressure (5 SNPs)^9^ and pulse pressure (2 SNPs)^9^. Of the 55 independent SNPs associated with C10AA (HMG CoA reductase inhibitors)-associated SNPs (P < 10^−8^/23), 19 SNPs have been reported to be significantly associated with low-density lipoprotein cholesterol (LDLC)^10^, supporting the known biological mechanism that statins are effective in lowering LDLC. However, for 3 medication-taking traits either small or no GWAS have been conducted for the medication-relevant indications, including A02B (drugs for peptic ulcer and gastro-oesophageal reflux disease), H03A (thyroid preparations) and N02BE (anilides).

**Figure 1.**
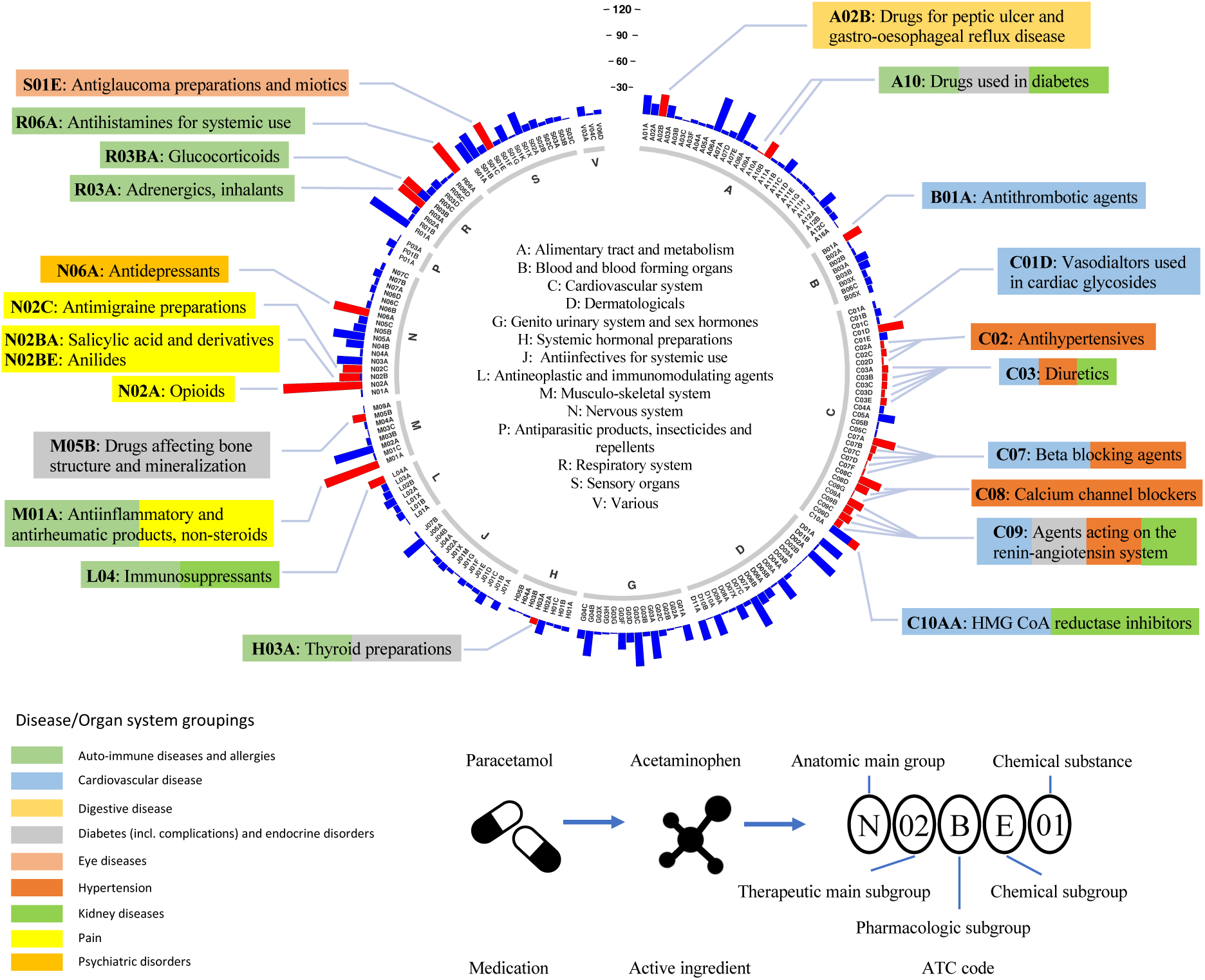
Distribution of 1,752 UKB medications at the first three ATC level. The inner ring corresponds to the 1^st^ level of the ATC code. The outer ring represents the first 3 level of the ATC code (184 subgroups). The length of the bar represents the number of classified UKB medications assigned to that subgroup (numbers of participants are shown in Figure 2). Red bars are the 23 medication-taking traits used in analyses (selected based on participant numbers). The 23 medication-taking traits are grouped into 9 diseases and organ system categories according to the main indications, which is highlighted using different colours (legend bottom left). The legend at the bottom right shows how ATC codes are assigned to each UKB medication.

**Figure 2.**
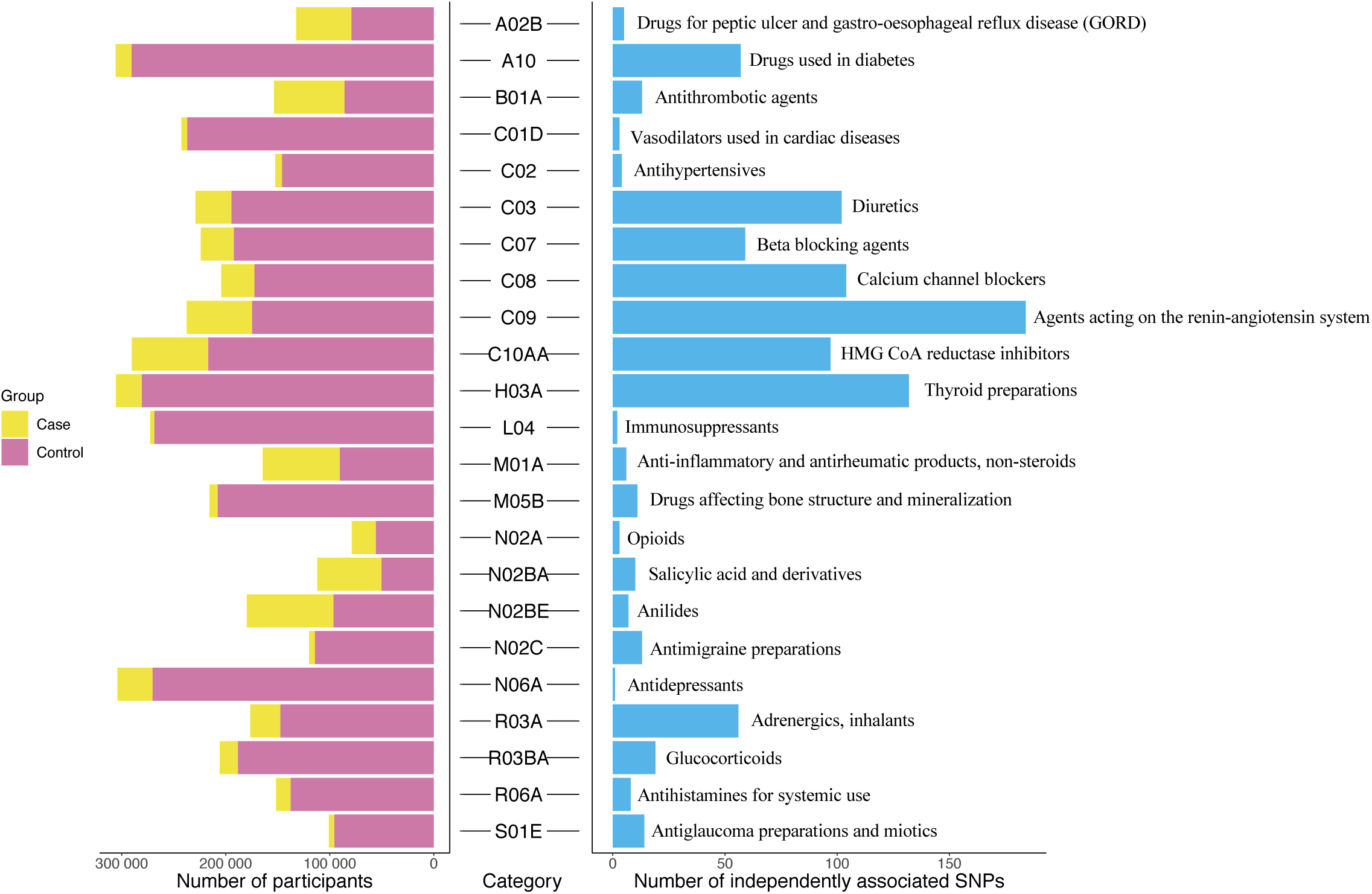
Summary of UKB medication-taking GWAS analyses. Text on the right side of each bar represents the meaning of each medication-taking ATC coded trait.

### Genetic predisposition to common disease predicts medication-taking

We undertook polygenic risk prediction analyses using GWAS summary statistics from 8 published disease/traits (**Table S5**) as discovery data to predict disease risk in 9 medication-taking phenotypes as target data. Participants in the UK Biobank with a high GRS for different diseases/traits have a higher odds of taking corresponding medications than those with a low GRS (**Figure 3**; **Table S6**). The top decile of individuals ranked on risk prediction for depression had an odds ratio (OR) of 1.7 in taking anti-depressants compared to the bottom decile. Similarly comparing top and bottom deciles, we find an OR of 3.1 for taking anti-diabetic medication (A10) for individuals ranked on genetic risk for type 2 diabetes and of 3.3 for taking immunosuppressants (L04) for individuals ranked on their genetic risk for rheumatoid arthritis (RA). The OR increased to 5.2 for taking L04 medications specific to RA (**Table S1**).

**Figure 3.**
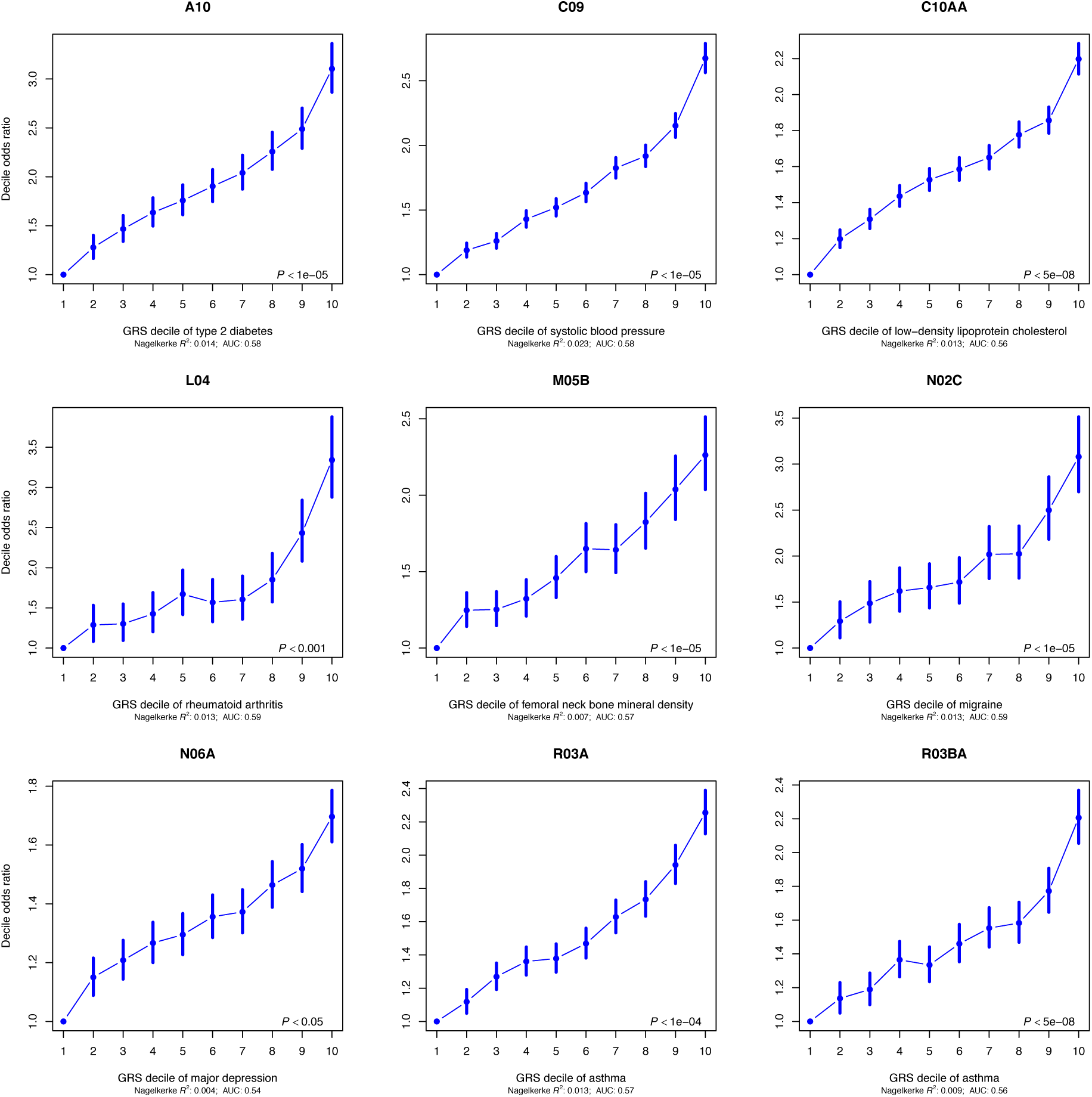
Odds ratio (OR) by genetic risk score (GRS) profile decile (1 = lowest, 10 = highest GRS), with OR reported relative to decile 1 as the reference. OR and 95% confidence intervals (blue bars) were estimated using logistic regression. The *P* value in the bottom right hand corner of each plot refers to the P-value threshold in the discovery sample used to generate the GRS. Note: An increased GRS of femoral neck bone mineral density implies a lower density.

### GWAS results and biological mechanisms

First, we estimated SNP-heritability of the 23 traits using linkage disequilibrium (LD) score regression^11^ (**Figure S6**; **Table S7**), all traits showed SNP-heritability (proportion of variance attributed to genome-wide SNPs) significantly different from zero to a maximum of 0.15 (s.e. 0.008) for N02A (opioid medications) on the estimated scale. Second, to identify medication-relevant tissue/cell types, we partitioned the SNP-heritability^12^ based on annotations of SNPs to genes, and genes to differential gene expression between tissues. Among the 23 medication-taking traits, 8 traits showed significantly enriched association with genes expressed in at least one tissue at a false discovery rate (FDR) < 5% (**Figure S7**). GWAS associations for thyroid preparations (H03A), immunosuppressants (L04), adrenergics inhalants (R03A), glucocorticoid (R06BA) and antihistamines for systemic use (R06A) were enriched in immune cell types. Those of opioid analgesics (N02A) were enriched in central nervous system tissues, such as limbic system, those of antimigraine preparations (N02C) were enriched in cardiovascular tissue, and those of drugs affecting bone structure and mineralization (M05B) were enriched in digestive cell type (**Table S8**).

Third, we investigated whether associations between SNPs and medication-taking traits were consistent with mediation through gene expression, based on associations between SNPs and gene expression (eQTLs). We identified 177 unique genes for which expression is significantly associated with 19 medication-taking categories (**Table S9**) using summary data-based Mendelian Randomization (SMR) analysis^13^. Gene-based association tests were conducted using MAGMA^14^ from the GWAS SNP results for each of the 23 medication-taking traits and a total of 1,841 significantly associated unique genes were identified (**Table S10**). To provide biological insights from the GWAS associated loci, we used the gene-based association test summary statistics to test for enrichment in 10,891 gene sets from MSigDB (v5.2)^15,16^. All 23 medication-taking traits were enriched in at least one gene set at FDR < 5% (**Table S11**). Several of the results showed plausible relevant biological mechanisms. For example, the genetic associations for taking A10 (drugs used in diabetes) were enriched for the glucose homeostasis gene set, those for taking C10AA (statins) were enriched in the cholesterol homeostasis gene set, C09 (agents acting on renin-angiotensin system) for cardiovascular-related gene sets, M05B (drugs affecting bone structure and mineralization) for skeletal system development, chondrocyte differentiation gene sets, N02A for gene sets of behavioural response to cocaine and neurogenesis and lastly H03A, L04, R03A, R03BA medications for immune-related gene sets. Interestingly, genes associated with taking A02B (drugs for peptic ulcer and gastro-oesophageal reflux disease) are enriched in gene sets of central nervous system neuron differentiation and of neurogenesis, highlighting the connection between gut and brain^17^.

### Linking genes associated with medication-taking to drug targets

Secondary analyses of GWAS results not only provide insights into the biological complexity of common diseases, but also offer opportunities relevant to drug development and repurposing^2,18,19^. To determine whether genes associated with medication-taking could provide clues relevant to drug target identification, we performed analyses using drug-target lists from Santos *et al.*^5^, ChEMBL^20^ and ClinicalTrial.gov (https://www.clinicaltrials.gov/) database as reference. First, for each UKB medication category, we investigated whether there are therapeutic-effect target genes for medications classified in that medication category; a total of 9 genes were identified (**Table S12**). For example, we find *HMGCR* (Entrez ID: 3156) is, as expected^21^, associated with taking C10AA medications (statins) and encodes the HMGCR protein which is targeted by medications from C10AA category. Second, we tested whether there are therapeutic-effect target genes for treating indications relevant to taking medications of each category; a total of 7 genes were identified (**Table S12**). *PCSK9* (Entrez ID: 255738) in our analyses is also associated with taking C10AA medications, and encodes the protein mediating lowering-cholesterol effect of evolocumab (ATC code: C10AX13) and alirocumab (ATC code: C10AX14). Third, we looked at whether there are therapeutic-effect target genes (ever or currently in clinical trial and not approved by FDA yet) for treating indications relevant to medications of each category; a total of 8 genes were identified (**Table S13**). For example, *TSLP* (Entrez ID: 85480) is associated with R03A (adrenergics), R03BA (glucocorticoids) and R06A (antihistamines) and also mediates the effect of tezepelumab for the treatment of uncontrolled asthma^22^. Hence, among our associated genes are 24 genes with some known evidence of therapeutic effect. Therefore, we anticipate that novel genes that are associated with medication may help to prioritise other putative therapies^23^. In **Table S14** we provide additional analyses for two genes, *IDE* and *AGT* that we believe merit further study for type 2 diabetes and C07/C09 related disorders, respectively.

### Shared genetic architecture between medication-taking traits and relevant complex traits

The genetic correlation (r_g_) between the 23 medication-taking traits and 21 traits/diseases (**Table S5**) related to them were calculated using bivariate LD score regression^24^. Many *r_g_* estimated were significantly different from zero. For example, body mass index, educational attainment (EA), former/current smoker and coronary artery disease were significantly correlated with most of the medication categories in expected directions. Major depression and neuroticism showed positive *r_g_* with A02B (gastro-oesophageal reflux drugs), suggesting a link between the brain and the digestive system. Type 2 diabetes showed correlations with taking medications C02, C03, C07~C09 and C10AA, implying a shared genetic architecture of type 2 diabetes, hypertension and hypercholesterolemia. The *r_g_* between B01A and N02BA show similar patterns of *r_g_* with other diseases/traits are similar to those for N02BA medications with other diseases/traits because the original medication aspirin (code number: 1140868226, 59150 individuals in our analysis) has multiple ATC codes (A01AD05, B01AC06 and N02BA01). Full results are presented in **Figure 4** and **Table S15**.

**Figure 4.**
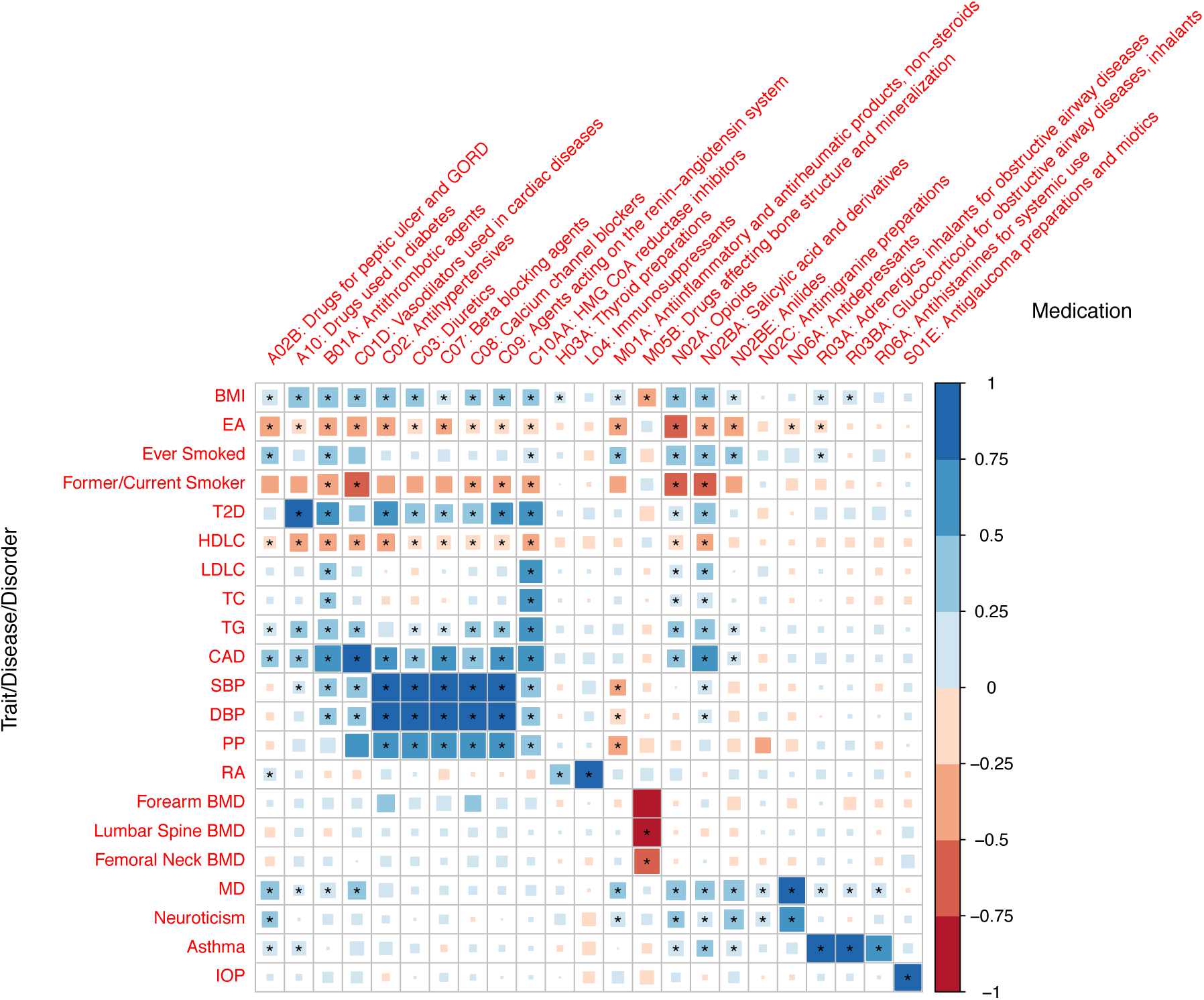
Genetic correlation of the 23 medication-taking traits and 21 diseases/traits related to them. Abbreviations: Body mass index (BMI), Education attainment (EA), Type 2 diabetes (T2D), High-density lipoprotein cholesterol (HDLC), Low-density lipoprotein cholesterol (LDLC), Total cholesterol (TC), Triglyceride (TG), Coronary artery disease (CAD), Systolic blood pressure (SBP), Diastolic blood pressure (DBP), Pulse pressure (PP), Rheumatoid arthritis (RA), Bone mineral density (BMD), Major depression (MD), Intraocular pressure (IOP).

### Putative causal relationship of diseases for using medication

It is reasonable to assume that having a disease is causal for taking the associated medication (rather than reverse causation). Therefore, we used Mendelian Randomisation (MR) in a proof-of-principle analysis to quantify causality. Independent SNPs (*P*-value < 5E-8) associated with 15 selected diseases/traits (**Table S5**) were used as instruments to evaluate putative causal relationships^25^ among these 15 diseases/traits and the 23 medication-taking traits (**Table S16 and Figure 5**). Increasing BMI increases the likelihood of taking A10, B01A, C01D, C02, C03, C07, C08, C09, C10AA, R03A medications, consistent with the role of BMI across diseases related to these medications^25^. The effect of obesity on bone health is controversial^26^. However, results from our analysis clearly show that increasing BMI decreases the likelihood of taking M05B (bone-associated) medications (OR 0.68 per SD of BMI). Major depression (MD) increases the likelihood of taking A02B medication (drugs for peptic ulcer and gastro-oesophageal reflux disease; 1.23-fold increase per SD in liability to MD), capturing a link between the brain and the digestive system. In addition to this, MD increases the likelihood of taking N02BE (1.23-fold increase per SD in liability to MD) medication, which is consistent with comorbidity of pain in some MD patients^27^.

**Figure 5.**
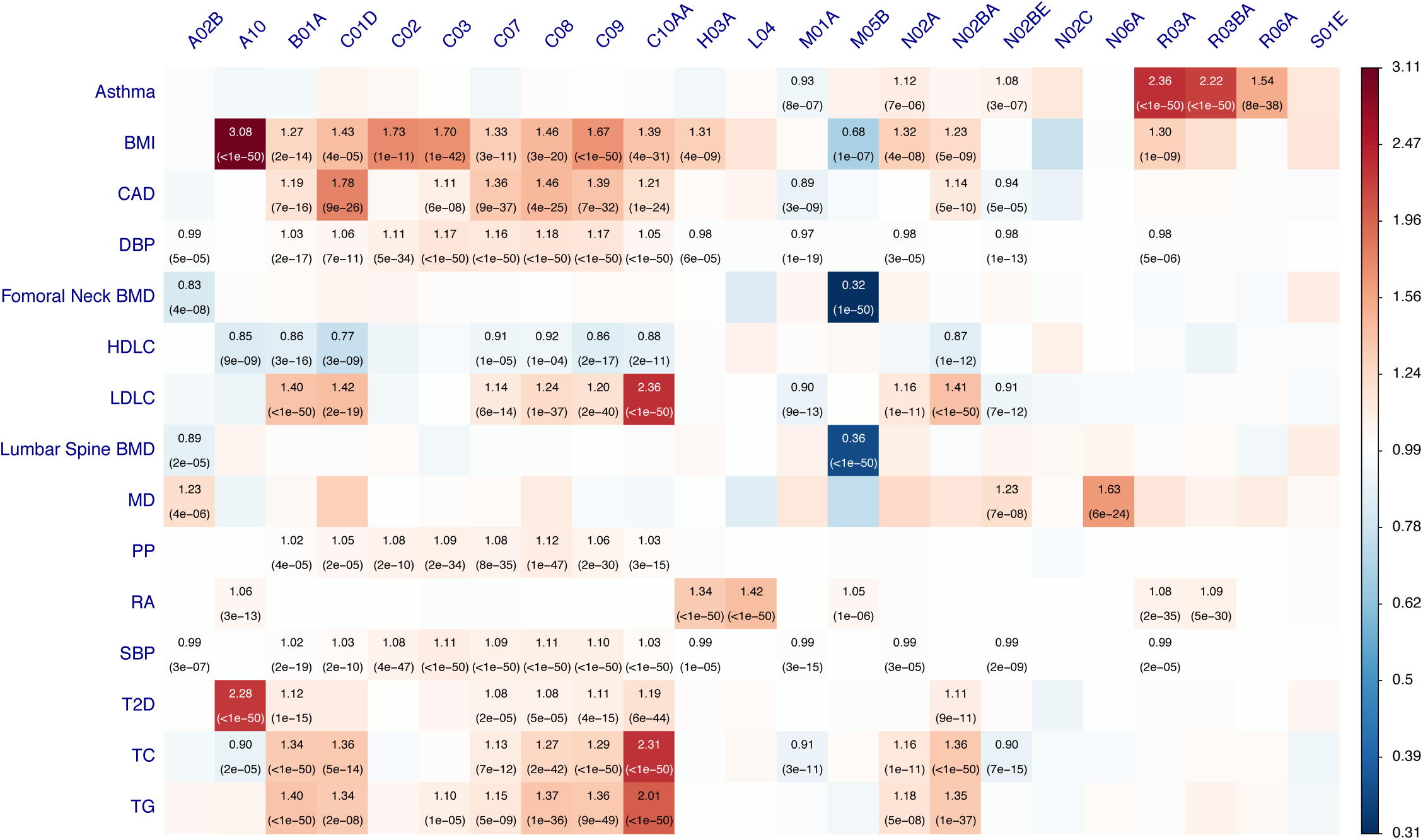
Mendelian Randomisation results using published SNPs associated with 15 diseases/traits as instrument. Rows are the exposure traits and columns are the outcome medication traits. Rows represent exposure and columns represent outcome. The significant effects after correcting for 345 tests (*P* value ≤ 1.4 × 10^−4^) are labelled with OR (P value). The OR is per SD in liability when the exposure is disease. Abbreviation: Body mass index (BMI), Coronary artery disease (CAD), Diastolic blood pressure (DBP), Bone mineral density (BMD), High-density lipoprotein cholesterol (HDLC), Low-density lipoprotein cholesterol (LDLC), Major depression (MD), Pulse pressure (PP), Rheumatoid arthritis (RA), Systolic blood pressure (SBP), Type 2 diabetes (T2D), Total cholesterol (TC), Triglyceride (TG).

## Discussion

To our knowledge, this is the first paper profiling genetic contributions to medication-use. Traditional GWAS identify DNA variants associated with disease, with a goal that these discoveries ultimately may open the door to new drug treatments. Here, we have taken the reverse approach, aiming to identify DNA variants associated with medication-taking, in recognition that underlying biology may contribute to the same medication being prescribed for several indications, and conversely that only some of those with a given diagnosis may take a particular medication. As expected, some of our results for medication-taking recapitulate GWAS results of the disease traits for which the medication is prescribed. However, we have also identified some novel associations that may be worthy of follow-up. We identified 505 linkage disequilibrium independent SNPs associated (P<1e-8/23) with different medication-taking traits. For some of our traits, large GWAS for the medication relevant indications have not been conducted, such as A02B (drugs for peptic ulcer and gastro-oesophageal reflux disease, 2 SNPs) and N02BE (anilides, 4 SNPs). Notably, 76 SNPs were associated with H03A (thyroid preparations - the main indication is hypothyroidism), only 11 of these loci have been previously reported to be associated with hypothyroidism. Conditional (mtCOJO) analysis suggested that these 76 SNPs associated with taking H03A medication are indeed associated with hypothyroidism. We showed that individuals with higher genetic risk of disease have higher likelihood to take relevant medications, for example, individuals with higher GRS for RA have an OR of 3.3 to take immunosuppressants compared with lower GRS individuals (**Figure 3**), thereby providing a proof-of-principle validation of precision medicine based upon risk prediction of common diseases, since individuals with high genetic risk of disease can be identified well before the onset of symptoms and the time of medication prescription.

To provide biological insight to the SNP associations for medication-taking^28^, we linked GWAS findings to relevant biological gene sets and drug target efficacy. These analyses generated a series of expected or plausible results, such as genes associated with taking A10 (drugs used in diabetes) enriched in gene sets for glucose homeostasis. Our analyses also generate new hypotheses; genes associated with taking N06A (antidepressants) showed enrichment in the gene set for the synthesis and secretion and diacylation of ghrelin, a gut-derived hormone^29^. Previous studies have described an antidepressant-like role of ghrelin^30,31^. This line of evidence suggests that testing a pharmacological effect of ghrelin on depression may be worthwhile. Although medication-associated genes overlapped with only a small proportion of current drug target genes, the framework of genetic association studies provides a potentially valuable resource for new drug target identification and prediction of unfavourable side effects^18^.

Comorbidity is commonly observed in clinical practice, which means the presence of additional diseases in relation to an index disease^32^. Results from genetic correlation and disease-medication (exposure-outcome) MR highlight potential shared aetiology, and may help explain medication use in clinical practice. Our analysis showed that major depression increased the likelihood of taking A02B (drugs for peptic ulcer and gastro-oesophageal reflux disease) and N02BE (anilides), the latter consistent with reports that antidepressant prescriptions are not only indicated for depression, but also for pain^33^.

There are a number of limitations in our study. First, although the medication-use data were obtained by trained nurses during interviews, the self-reported nature may limit the accuracy of information. Second, the ambiguous names of medications may limit the accurate classification of medications. The reasons (e.g. disease diagnosis) for taking medication were not recorded and hence not available for further analysis, nor were duration and dosage of medications. Third, our findings are specific to the UK biobank participants, which are recognized to be a non-random sample of the UK population. Fourth, the medication-taking in UK biobank participants may be more representative of medication-taking in the UK and may not translate to other populations and different health systems.

In summary, we identified 505 independent loci associated with different medication-use in 318,177 individuals from UKB, with implications for biological mechanisms, drug target identification and precision medicine for common disease.

## Methods

### Medication data

We used self-report data of regular medication (prescription and over-the-counter) and health supplements taken weekly, monthly or three-monthly from participants in the United Kingdom Biobank (UKB) study (http://www.ukbiobank.ac.uk)^34^, mainly aged 37–73 years when recruited between 2006 and 2010. Medication and health supplements data were coded and manually mapped to their corresponding active ingredients and then to their Anatomical Therapeutic Chemical (ATC) Classification System^5^ codes (**Table S1**). In total, medications were classified into 1,752 categories, collapsing to 184 subgroups according to the first three ATC levels (**Figure 1, S1**). 23 of these medication subgroups (based on participant numbers) were selected for analysis. Detailed methods are provided in the **Supplementary Appendix**.

### Genome-wide association study design and statistical analysis

Analyses used genome-wide genotypes for 318,177 participants of white European descent (**Figure S2**). 23 medication-taking case groups and their corresponding control groups were generated. Case groups were defined as those taking medications classified at the same ATC level. Control groups comprised participants taking neither the case medication nor similar medications. Similar medications were defined at those sharing the first two ATC levels as the case medication or medications containing the case medication active ingredients. Following standard quality control and genotype imputation methods (see **Supplementary Appendix**), 7,288,503 SNPs with minor allele frequency (MAF) > 0.01 were used in analyses. Case-control genome-wide association analyses were conducted using BOLT-LMM^35^ with age, sex, assessment centre and 20 genetic principal components fitted as covariates. Conditional analyses tested if SNPs associated with taking medications have been previously linked to their corresponding medication-specific related indications/traits^36,37^. Multi-trait-based conditional and joint (mtCOJO) analyses tested if medication-taking associated SNPs are also associated with their relevant main indications in UKB^25^.

Genetic risk score (GRS) for UKB individuals were generated for 8 diseases using SNP effect size estimates from published GWAS summary statistics (discovery sample data) (**Table S2**). These GRS were used to predict the medication use traits related to these diseases (asthma mapped to two medication use traits). Selection of the discovery samples data was based on relationship to the medication-taking traits, availability of GWAS summary statistics, cohort ancestry and no sample overlap with UKB. GRS were generated for a range of discovery data association *P* value thresholds (5×10^−8^, 1 × 10^−5^, 1 × 10^−4^, 0.001, 0.01, 0.05, 0.1, 0.5). The GRS were evaluated as medication-taking odds ratio for each GRS decile (relative to the 1^st^ decile).

LD score regression^11,24^ was used to estimate the proportion of variance attributable to genome-wide SNPs 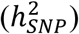 and to quantify genetic sharing at common variants across the 23 medication-taking traits and other traits. LD score regression for cell type specific analysis^12^ was applied to test the 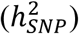 enrichment in different tissues for each of the 23 medication-taking traits. Gene expression data of 205 tissues (53 from GTEx^38^ and 152 from other sources^39,40^) were used for analyses. Summary-data-based Mendelian Randomization (SMR)^13^ was used to integrate our trait association with blood expression quantitative trait loci (eQTL, i.e., SNP-gene expression association) data^41^. Gene-based association analyses were conducted using MAGMA (v1.06)^14^ to identify genes associated with different medication-taking traits. Gene sets association analyses were conducted using MAGMA (v1.06)^14^ with curated gene sets (c2.all) and gene ontology sets (c5.bp, c5.cc, c5.mf) from MSigDB (v5.2)^15^,^16^.

Mendelian Randomization (MR) was used to investigate a putative causal relationship between the 23 medication-taking traits and 15 significantly correlated traits (selected from **Table S2**), using Generalized Summary-data-based MR (GSMR)^25^. We required that all analyses had at least 7 genome-wide significant loci to use as MR instruments; the median number of SNP instruments was 65.

### Analyses linking GWAS results to drugs and disease

To check whether associated genes from MAGMA and SMR encode effect-mediating targets for FDA-approved medications or corresponding indications, we used information from Santos *et al.^5^*, based on medication approved by the FDA before June 2015. For those approved later, we used the ChEMBL database^20^. To check whether associated genes encode trait-relevant effect-mediating targets for drugs in clinical trial, we used ClinicalTrial.gov (https://www.clinicaltrials.gov/). The CLUE Touchstone tool (https://clue.io/)^42^ was used to check the correlation between signatures of drugs and knocking down a gene.

### URLs

UK Biobank: http://www.ukbiobank.ac.uk; ClinicalTrial.gov: https://www.clinicaltrials.gov/; CLUE Touchstone tool: https://clue.io/.

## Acknowledgments

We thank members of The University of Queensland Program in Complex Trait Genomics group. This research was supported by the Australian National Health and Medical Research Council (Grant Number: 1113400, 1078901, 1078037). Yeda Wu is supported by the F.G. Meade Scholarship and UQ Research Training Scholarship from the University of Queensland. This study makes use of data from UK Biobank (Project ID: 12514) and we thank the UK Biobank participants and the UK Biobank team for generating an important research resource.

## Author contributions

P.M.V., N.R.W. and Y.W conceived and designed the experiment. Y.W. performed the analysis with assistance and guidance from E.M.B., Z.Z., K.E.K., J.Y., Z.Z., K.E.K. and L.Y. contributed to data quality of UKB data. Y.W., P.M.V. and N.R.W. wrote the manuscript with the participation of all authors.

