## Supplementary Appendix for "Genetic analyses of medication-use and implications for precision medicine"

This appendix has been provided by the authors to give readers additional information about their work. Supplement to: Wu Y, Byrne E, Zheng Z, et al. Genetic analyses of medication use and implications for precision medicine.

### Supplementary text

#### United Kingdom Biobank (UKB)

The study design and sample characteristics of the United Kingdom Biobank (UKB) (http://www.ukbiobank.ac.uk), a major population-based longitudinal study, have been extensively described elsewhere[1]. The UKB was approved by the National Research Ethics Service Committee North West Multi-Centre Haydock and all participants provided written informed consent to participate in the study. Briefly, initial data from more than 500,000 individuals aged 37-73 years were collected between 2006 and 2010, with first-repeat data (revisit) collected on approximately 20,000 individuals from 2012 to 2013 and second-repeat data (imaging visit) collected from approximately 22,000 individuals since 2014 (Downloaded on March 2017).

#### UKB medication classification

Self-reported regular medication and health supplements taken weekly, monthly or three monthly were recorded. Duration and dosage of the medication records were not collected[2]. Medication and health supplements data (Data Field: 20003) were coded using 6,745 categories (Data coding 4). 1,809 of 6,745 categories were for medications taken by at least 10 participants. These 1,809 categories were manually mapped to their corresponding active ingredients using online information (mainly Electronic Medicines Compendium (https://www.medicines.org.uk/emc), Drug (https://www.drugs.com/), and NetDoctor (https://www.netdoctor.co.uk/)), and were classified using the Anatomical Therapeutic Chemical (ATC) Classification System[3]. Categories named by their active ingredient(s) were directly mapped to the ATC code. Categories named by brand name were first mapped to their active ingredient(s) and then further mapped to ATC code according to the dose and administration route if available. Some categories were ambiguous or could not be mapped to an ATC code, leaving 1,752 categories which grouped in to 184 subgroups according to the first three ATC level (**Figure 1** and **S1**). **Table S1** provides the active ingredient(s) and ATC code information for the 1,752 categories.

#### UKB medication taking demographics

There were 502,616 participants (~54% females) with medication records (~72.5% with non-blank medication information) at their first visit UKB assessment. The mean age for the participants when attending assessment centre was 56.53 (s.e. 8.09) years and the mean body mass index (BMI) for participants was 27.43 (s.e. 4.80). The percentage of participants taking medication increased with age. The percentage of female participants was always higher than that of males, across all age groups, but the percentage of males taking medication increased sharply from ~50% at 40 years old to ~85% at 70 (**Figure S2A**). The number of medications per participant increases with age (**Figure S2B**).

#### UK Biobank genotyping, quality control and participants selection

Genotyping details for UKB participants have been reported previously[4]. Briefly, 49,950 participants were genotyped using the UK BiLEVE Axiom Array and 438,427 participants were genotyped using UK Biobank Axiom Array. The Haplotype Reference Consortium

(HRC) and UK10K was the imputation reference sample. A European subset (456,414 participants) were identified by projecting the UK Biobank participants onto the 1000 Genome Project principal components coordination. Genotype probabilities were converted to hard-call genotypes using PLINK2 (--hard-call 0.1) and single nucleotide polymorphisms (SNPs) with minor allele count <5, Hardy-Weinberg equilibrium test *P*-value < $1\times10^{-6}$, missing genotype rate > 0.05, or imputation info score < 0.3 were excluded. Following the phenotype extraction pipeline for UKB participants provided in **Figure S3**, 318,177 participants of European ancestry with both genotype and medication records available were selected for further analysis.

### Case-control genome wide association study (GWAS) designs

Case group and control group were generated according to case medications, similar medications and control medications. Medications with the same ATC level (at the first two, the first three and the first four level) were defined as case medications and those taking case medications were assigned to the corresponding case group. Medications of which the first two ATC level is the same as that of the case medication active ingredients or medications containing case medication active ingredients were defined as similar medications. After excluding participants taking both case medications and similar medications, the remaining participants were assigned to corresponding control group (those taking "99999" category were removed). A total of 23 case-control medication category traits were selected for analysis. Case-control GWAS analyses were conducted using BOLT-LMM[5] with age, sex, assessment centre and 20 genetic principal components fitted as covariates. 543,919 SNPs generated by LD pruning ($r^2 < 0.9$) from Hapmap3 SNPs were used to control for population structure and polygenic effects. The effect size (betas) and standard error (se) from BOLT-LMM on the observed 0-1 scale were transformed to odds ratio (OR) and corresponding standard error (SE) using log OR = β / (P* (1 - P)) and SE = se / (P * (1 - P)), where β = linear regression coefficient, se = standard error from BOLT-LMM and P = case fraction. 7,288,503 SNPs with minor allele frequency (MAF) > 0.01 were analysed. Quasi-independent trait-associated regions were generated through linkage disequilibrium (LD) clumping retaining the most associated SNP in each region (PLINK (v1.90b)[6,7] --clump-p1 5e-8 --clump-p2 5e-8 --clump-r2 0.01 --clump-kb 1000). If associated, the MHC region (25Mb-34Mb) was considered as a single locus represented by its most associated SNP. To explore how many SNPs associated with taking medications have been previously linked to their corresponding medication-specific related indications/traits, GCTA (v1.91) was used to perform analyses (--cojo-cond) of 10 medication GWAS summary statistics, conditioning on the given lists of SNPs associated with relevant indications/traits[8,9]. The GWAS Catalog[10] was used to search published GWASs on relevant indications/traits, with studies selected based on number of independent SNPs reported. To check whether medication-taking associated SNPs were also associated with the main indications for that medication category, GCTA (v1.91) was used to perform analyses (--mtcojo-file) of the 10 medication GWAS summary statistics, conditioning on the related main indications GWAS summary statistics in UKB. The indication phenotype were generated using self-reported non-cancer illness code (Data Field: 20002), main ICD10 diagnoses (Data Field: 41202) and secondary ICD10 diagnoses (Data Field: 41204).

### Genetic risk score (GRS) prediction

Of the 23 medication-taking traits, related published GWAS summary statistics (discovery data) were available for 9 of these medication-taking traits (target data), based on 8 discovery

GWAS studies (***Table S5***). Discovery data were selected as traits related to target data phenotypes, cohort ancestry and with no sample overlap with UKB. The discovery data SNPs were matched with the target data SNPs, then LD pruned and “clumped”, discarding variants within 1,000 kb of, and in $r^2 \geq 0.1$ with, another (more significant) marker using unrelated European participants in UKB as an LD reference. Genetic risk scores of target sample individuals were generated for a range of discovery data association *P* value thresholds ($5\times10^{-8}$, $1\times10^{-5}$, $1\times10^{-4}$, 0.001, 0.01, 0.05, 0.1, 0.5). For each discovery-target pair, four outcome variables were calculated. 1) The *P* value of case-control GRS difference was calculated by logistic regression. 2) The proportion of variance explained (*Nagelkerke's* $R^2$) was calculated by comparison of a full model (phenotype ~ GRS) with a null model (phenotype ~ 1). 3) Area under the receiver operator characteristic curve using R package pROC[38], which can be interpreted as the probability of ranking a randomly chosen case higher than a randomly chosen control. 4) Odds ratio and 95% confidential interval for the 2nd to 10th GRS deciles group compared with 1st decile. GRS were converted to deciles from lowest (1) to highest (10) GRS.

### Linkage disequilibrium (LD) score regression

Heritability attributable to genome-wide SNPs estimated on the sample scale ($h^2_{SNP}$) were estimated using LD score regression[11] from the GWAS summary statistics of 23 medication-taking traits. To evaluate the extent of shared common variant genetic architectures between the 23 medication-taking traits and a range of human traits, disorders and diseases[12-24] (***Table S5***), the bivariate genetic correlations[25] attributable to genome-wide SNPs ($r_g$) were also calculated using LD score regression.

### Linking GWAS findings to gene expression

LD score regression for cell type specific analysis[26] was applied to test the enrichment heritability in different tissues for each of the 23 medication-taking traits. Gene expression data of 205 tissues (53 from GTEx[27] and 152 from Franke lab[28,29]) were used for analysis. Summary-data-based Mendelian Randomization (SMR)[30] was used to identify the causal relationship between gene expression and trait. Westra expression quantitative trait loci (eQTL) data[31] were used in the SMR analysis.

### Gene-based association and gene sets analysis

MAGMA (v1.06)[32] was used to compute mean association *P* values for a gene-based test. SNPs with MAF > 0.01 from 10,000 random sampled unrelated UKB European-ancestry individuals were used as the LD reference. The window size used was 35 kilobase (kb) upstream and 10 kb downstream to include regulatory elements. The SNPs were mapped to a total of 18,348 genes for each trait using gene locations (build 37) file. For gene sets analysis, curated gene sets (c2.all) and gene ontology sets (c5.bp, c5.cc, c5.mf) from MSigDB (v5.2)[33,34] were tested for each of the 23 traits. Competitive test *P* values for each gene set were computed; correcting for gene size, density, minor allele count and gene-gene correlations[32]. We generated FDR-adjusted *P* values for biological pathways using Benjamini and Hochberg's method to account for multiple testing[35].

### Linking GWA findings to drug target

To check whether a gene encodes an effect-mediating target for FDA-approved medications or corresponding indications, we adopted the supplementary tables from Santos *et al.*[3]. However, this table contains medication approved by FDA before June 2015. For those approved later, we used the ChEMBL database[36]. To check whether gene encodes effect-mediating target for drug in clinical trial corresponding to associated medications' indication, we further used ClinicalTrial.gov (https://www.clinicaltrials.gov/). To check the correlation between signatures of drugs and knocking down a gene, the Touchstone tool from CLUE website (https://clue.io/)[37] was used.

### Mendelian randomization (MR)

MR was used to investigate the causal relationship between the 23 medication-taking traits and other significantly correlated traits. The correlated traits were selected from ***Table S5***. We required that the data samples all had ≥7 genome-wide significant loci to use as MR instruments; the median number of SNP instruments was 65. 15 correlated traits were used to conduct MR analysis using Generalized Summary-data-based MR (GSMR) [39], which includes a heterogeneity test to exclude highly pleiotropic loci. The other parameters were set as default in GCTA-GSMR.

### Supplementary figures

#### Figure S1. Analysis pipeline of UKB medication

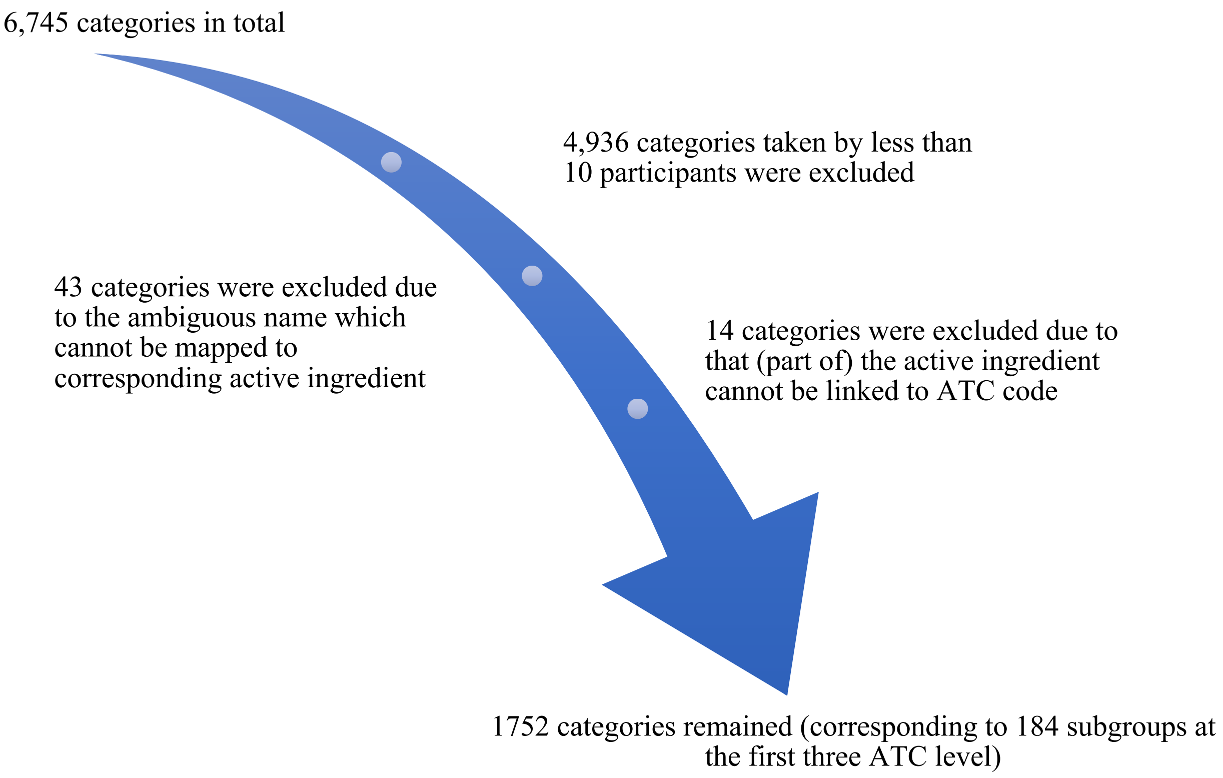

**Figure S2. Demographic statistics for participants who completed self-report medication questionnaires at first visit in Biobank**

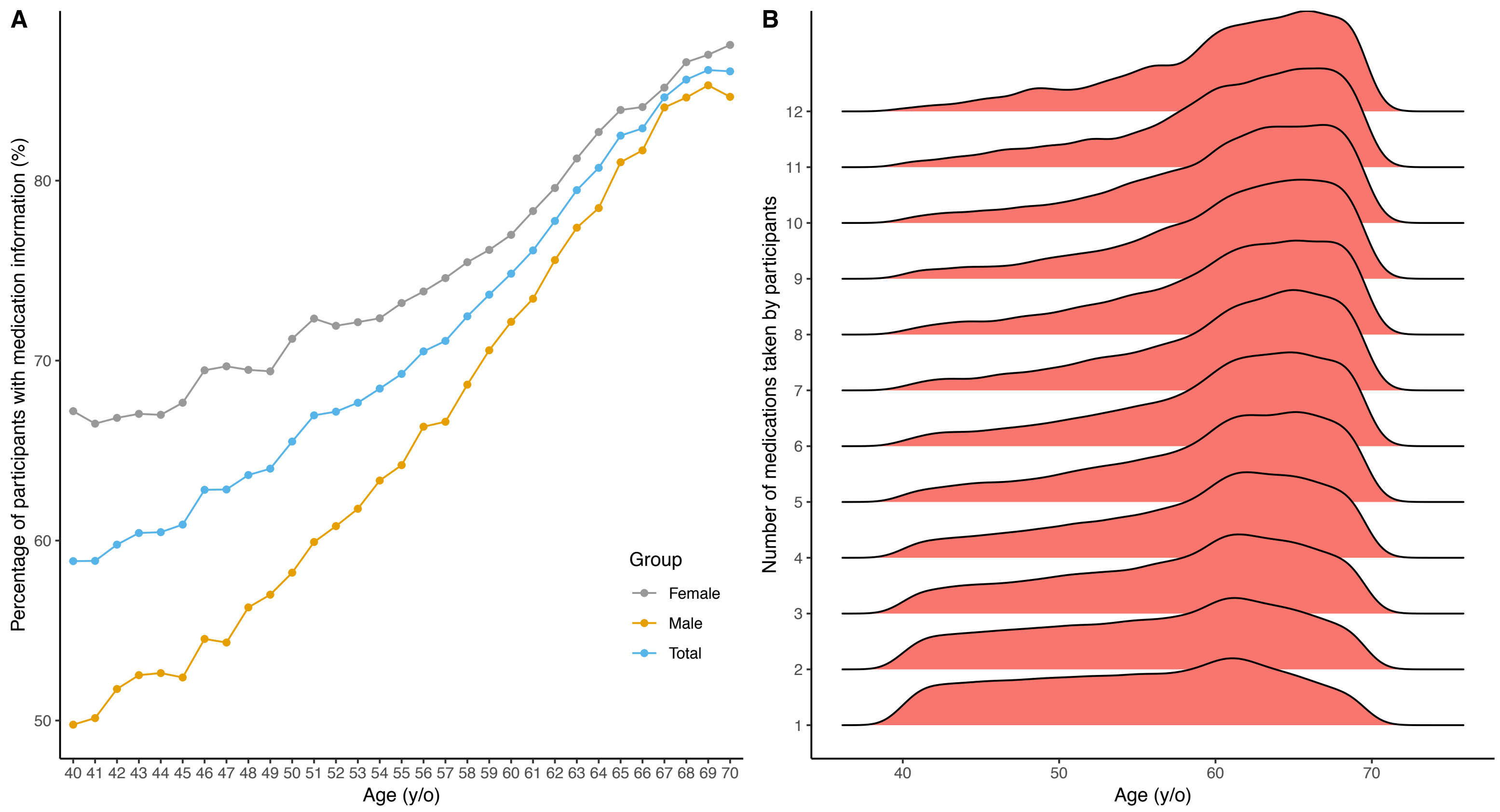

Figure S3A. Percentage of participants with medication information at different ages.
Figure S3B. Age distribution of participants stratified by the number of medications taken.

**Figure S3. Phenotype extraction pipeline for UKB participants**

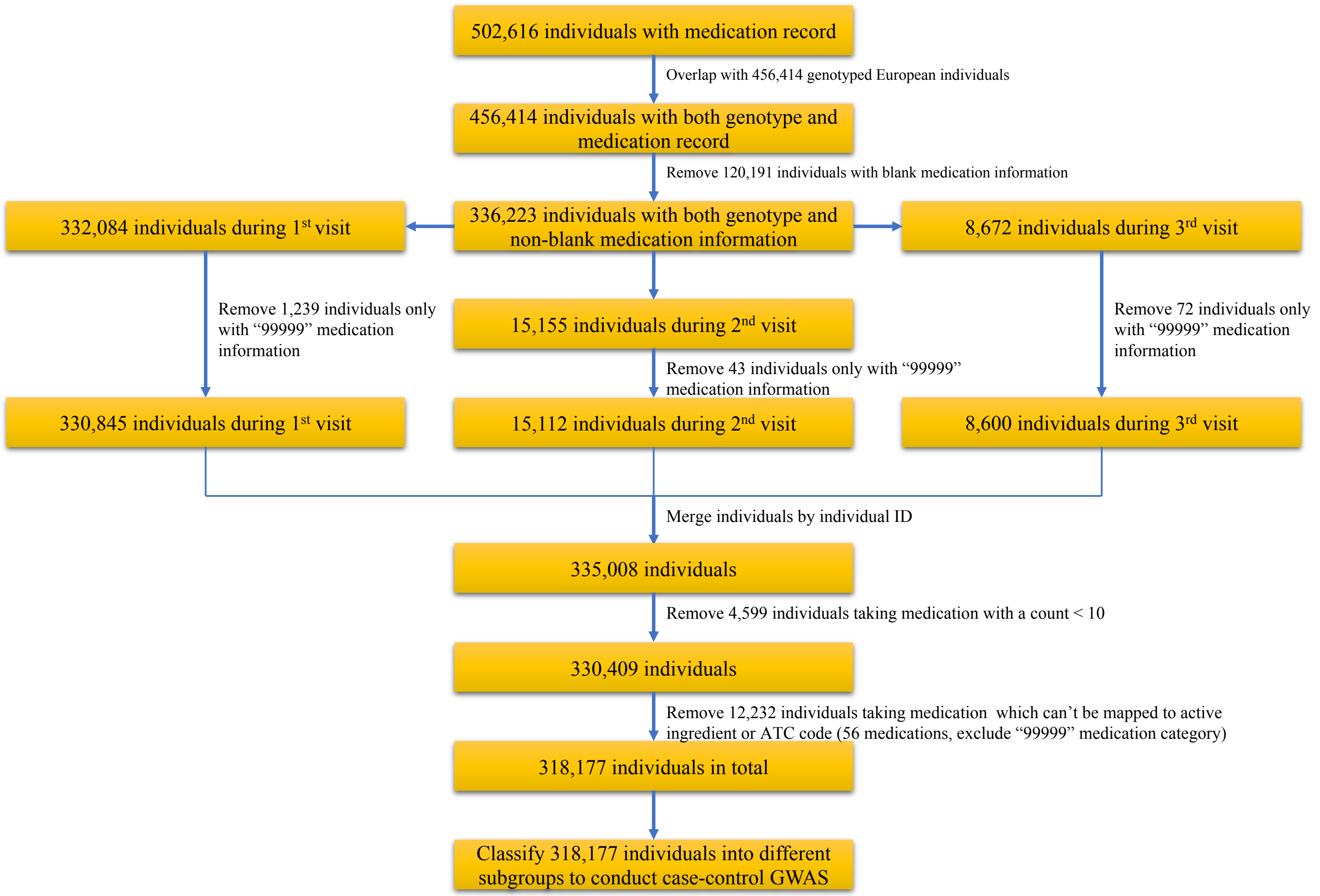

**Figure S4. Full analysis workflow of the study**

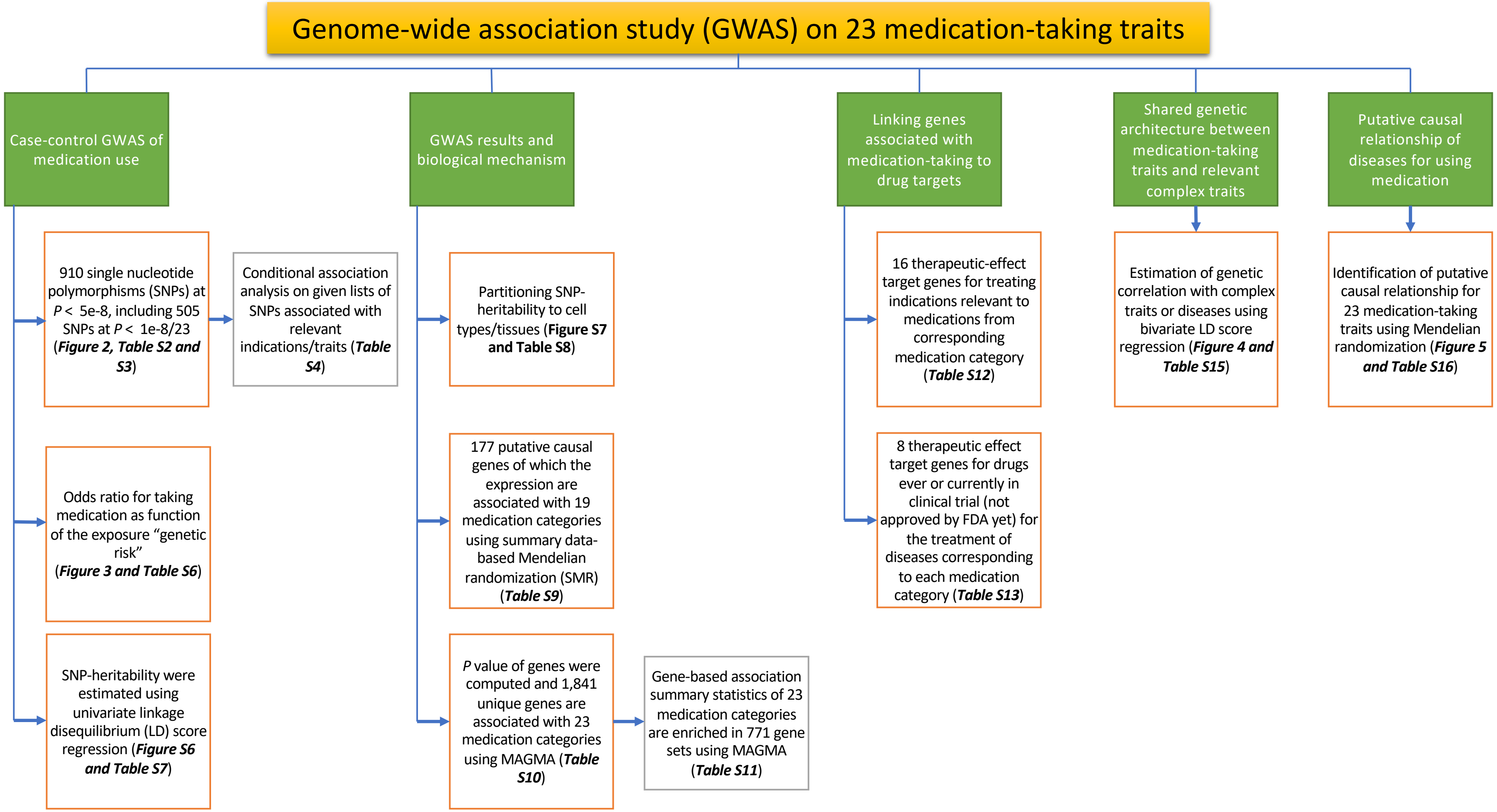

### Figure S5. Manhattan plot for 23 medication-taking traits

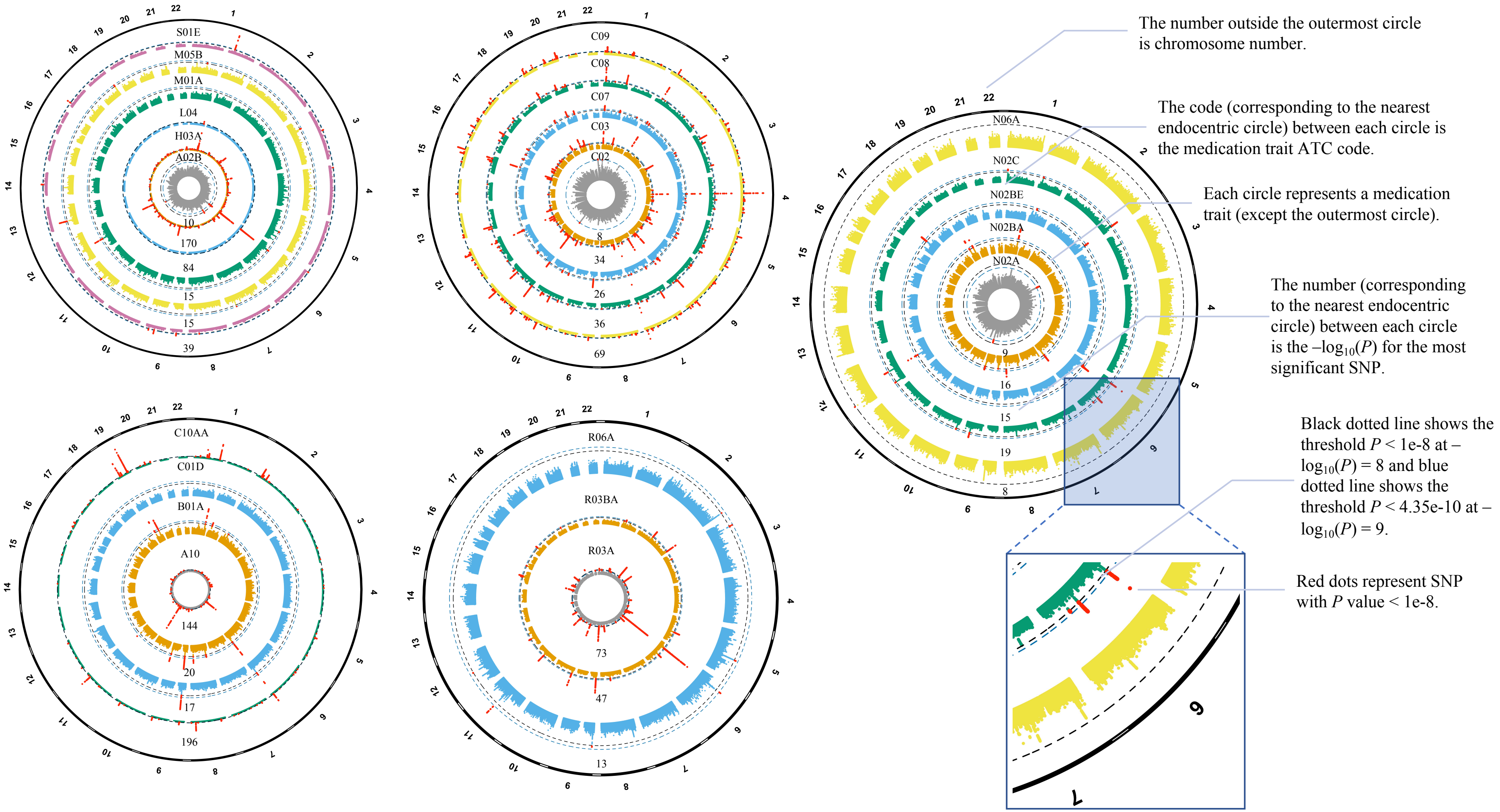

**Figure S6. The SNP-heritability of medication-taking trait**

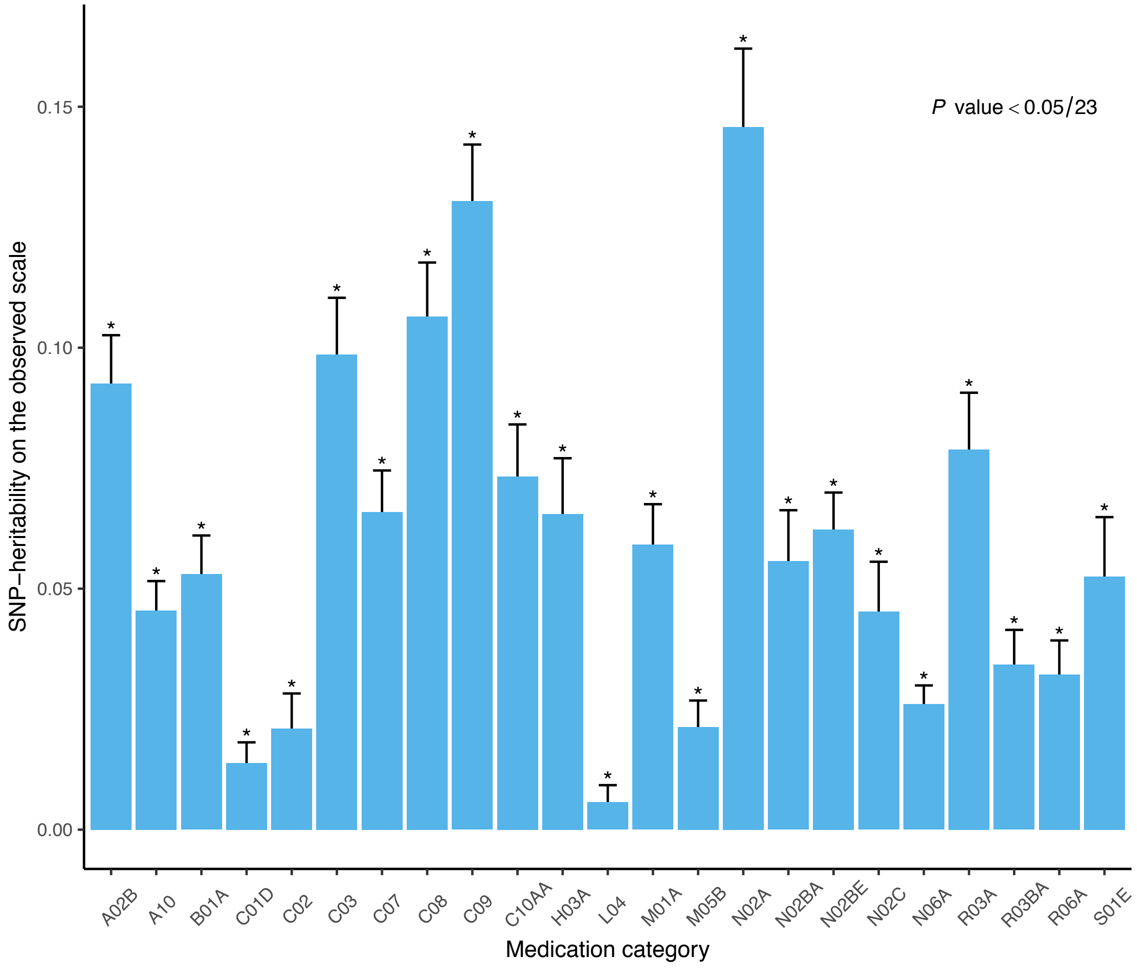

### Figure S7. Results of the multiple-tissue analysis for 8 medication-taking traits

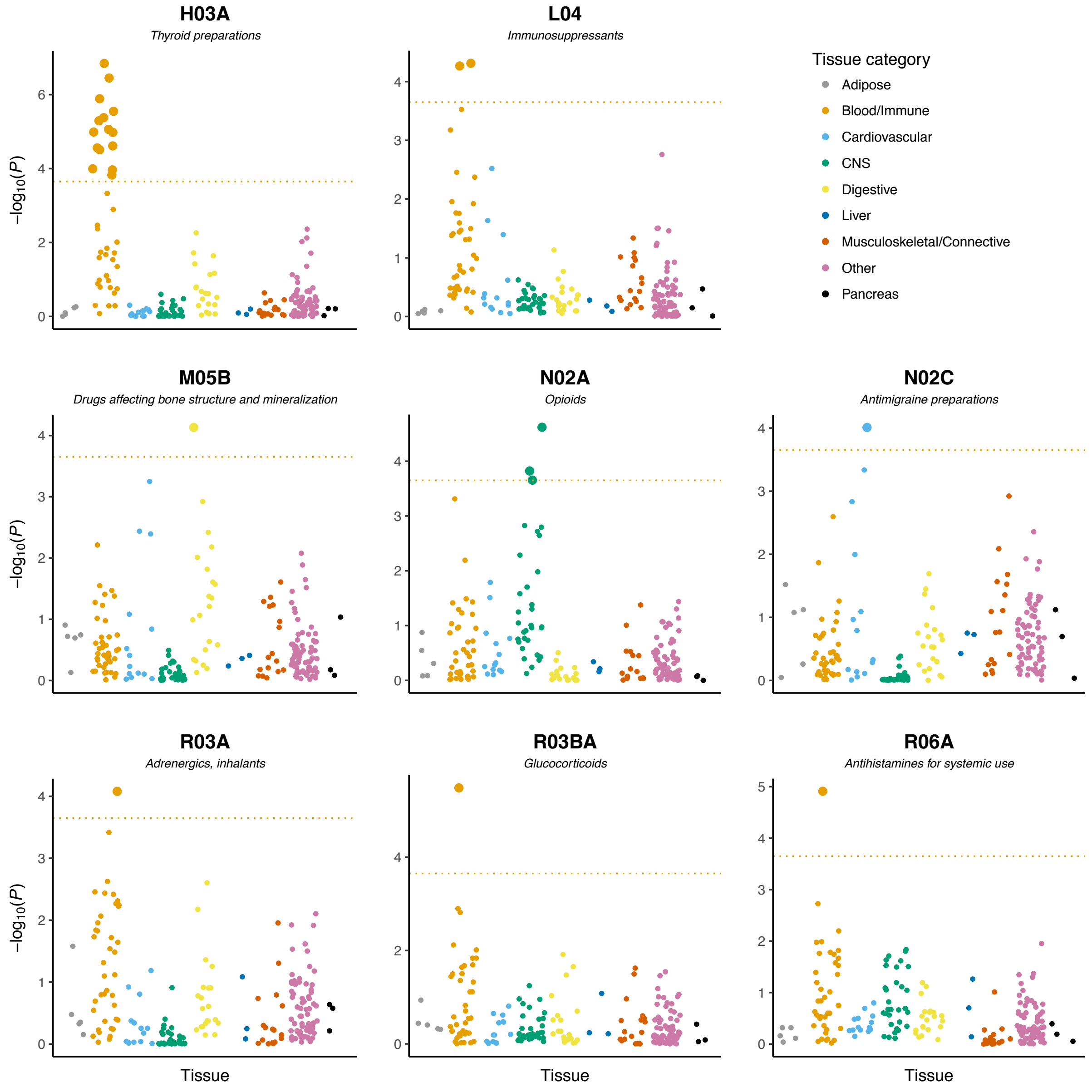

Each dot represents a tissue or cell type. The dotted line shows the threshold of FDR < 5% at -$\log_{10}(P)$ = 3.81. Large dots pass the threshold.
